# The Invariant Nature of a Morphological Character and Character State: Insights from Gene Regulatory Networks

**DOI:** 10.1101/420471

**Authors:** Sergei Tarasov

## Abstract

What constitutes a morphological character versus character state has been long discussed in the systematics literature but the consensus on this issue is still missing. Different methods of classifying organismal features into characters and character states can dramatically affect the results of phylogenetic analyses. Here, I show that the modular structure of the gene regulatory network (GRN) underlying trait development, and the hierarchical nature of GRN evolution, essentially remove the distinction between morphological character and character state, thus endowing the character and character state with an invariant property with respect to each other. This property allows representing the states of one character as several individual characters and vice versa. In practice, this means that a phenotype can be encoded using a set of characters or just one complex character with numerous states. The representation of a phenotype using one complex character requires a selection of an appropriate penalty for the state transitions.

## INTRODUCTION

The distinction between character and character state has long been debated in the phylogenetic literature (reviewed in [1, 2]), making it difficult for researchers to know which concept of the character to use. Despite the numerous definitions, there is no consensus among researchers regarding what constitutes a character versus character state. One group of the researchers suggest that both concepts are the same [3–5]. However, others argue that character and character state are essentially different [6–12]. A recent view supporting the distinction treats character states as mutually exclusive observations that have to be combined together into a character to perform an analysis [2]. Under this view, characters are classified into two fundamental categories: the neomorphic, referring to the birth or loss of a morphological feature, and the transformational, referring to transformation of the feature from one state to another. Yet another view provides a similar distinction, but places the character versus character state debate in the context of evolutionary development (evo-devo). Here, each character is determined by a character-specific gene regulatory network (GRN, see Box 1) that is called the character identity network [11, 12]. The character identity network triggers the realization of downstream GRNs, which determine character states. The different ways of classifying organismal features into characters and character states can affect the results of phylogenetic analyses [13]. This makes it imperative to find a unified and rigorous approach to deal with the two concepts. Below, I will show that the equivalence of character and character state can be directly derived from the structure and appropriate representation of the developmental GRNs that underlie the evolution of the morphological traits. This equivalence indicates that states of the same character can be represented as separate characters and, at the same time, separate characters can be combined into one single character. Thus, the character and character state remain conceptually unchanged, and are thereby invariant with respect to each other, regardless of the way of classifying organismal features into characters and states.

### Mechanisms of GRN evolution

Numerous evo-devo studies shed light on the mechanisms that drive evolution of GRNs and their corresponding traits across species. The evolution of a GRN, to a certain extent, is dictated by its modular structure (Box 1) where each GRN module (including the GRN of an entire organism) consists of the smaller GRN submodules which regulate the expression of genes during development thus giving rise to observed phenotypic variations [14–21].

**BOX 1.**
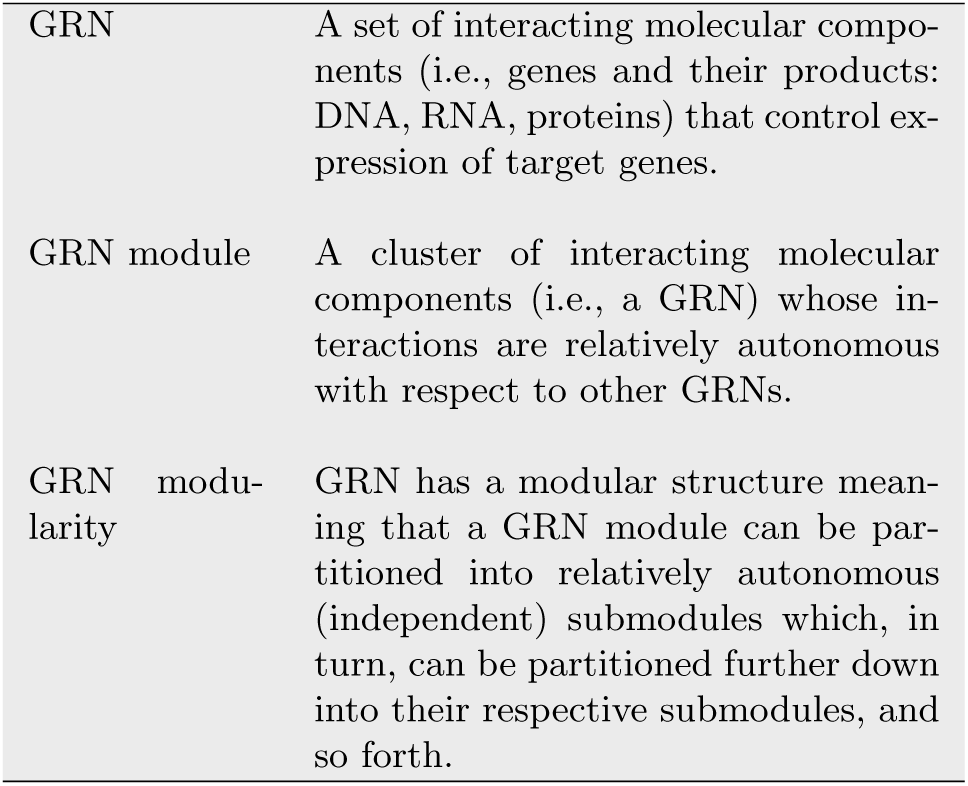
Denitions of the key terms.

Therefore, the evolution of morphological features is the evolution of the GRN modules. In general, morphological features evolve largely due to mutations in the regulatory loci (e.g., cis-regulatory elements), which control the expression of functionally conserved proteins [22]. The core mechanisms of GRN evolution include the elementary genetic processes of mutation and recombination: nucleotide substitutions within a sequence, site insertion/deletion, replication and rearrangements of sequence fragments within the genome. All mechanisms mediating GRN evolution can be classified into the following four categories (Fig. 1).

**Figure 1.**
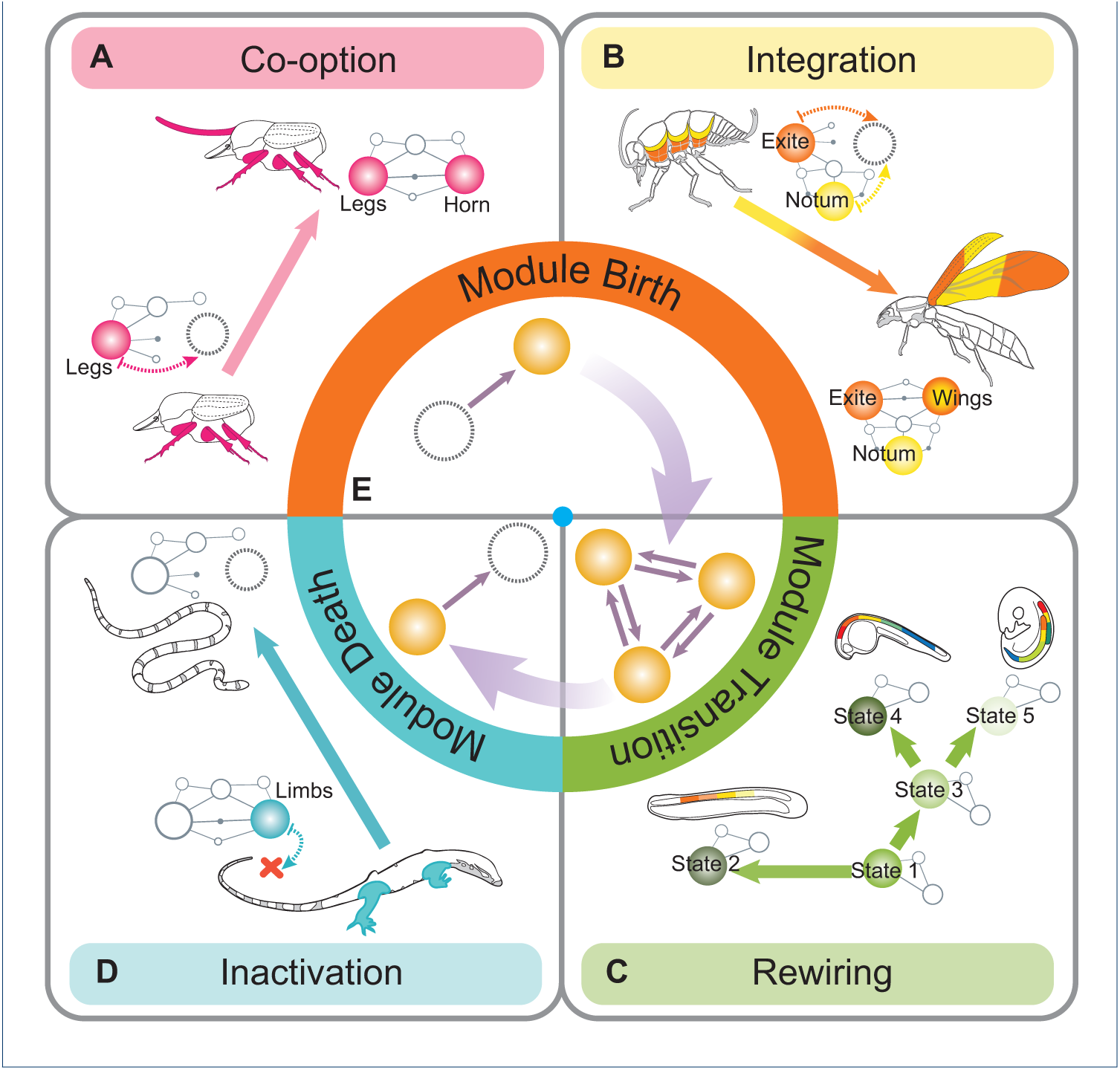
The mechanisms of GRN evolution. The colored balls are the GRN modules, the dashed balls indicate the absence of a module. **(A)** The mechanism of module co-option resulting in the formation of horns in dung beetles. The pre-existing module responsible for the development of beetle appendages (Legs, the rose ball) is co-opted into the prothorax forming the pronotal outgrowth (Horn, the rose ball). **(B)** Module integration resulting in the formation of the insect wings. The wings (the orange-yellow ball) evolved by merging two GRN modules; these two modules control development of the leg exites (the orange ball) and the lateral extension of notum (the yellow ball); the insect drawings were redrawn from [32]. **(C)** Module rewiring; the rewiring of modules within the Hox gene cluster generated the diversity of animal body plans during an early stage of metazoan evolution. The rewiring is exemplified as the transition between different states of the *Hox* genes cluster (the green balls). The animal body plans were redrawn from [37]. **(D)** Module inactivation resulting in loss of legs in snakes. A mutation in ZPA enhancer switches-off the formation of a downstream GRN module that controls limb development (Limbs, the blue ball). **(E)** The full evolutionary path of a GRN module that shows module birth, module transition, and module death.

### 1. Module co-option (Fig. 1A)

This mechanism assumes recruitment of a pre-existing module into a novel place in the body. Such process can be accompanied by some rewiring of the co-opted module, and exhibits a widespread mechanism that produces novel traits and serial homologs [12, 14, 19, 20, 23–27]. For example, the males of dung beetles are famous for the shapes of their cephalic and thoracic horns, which they use in combat for access to females. It was shown [28] that these horns evolved by the co-option of GRN that participates in the development of beetle appendages into a new body region (Fig. 1A).

### 2. Module integration (Fig. 1B)

This mechanism implies emergence of a new GRN module by integration of pre-existing modules [29]. The origin of insect wings has been debated for a long time, with two alternative hypotheses: one stating that wings evolved from the leg exites [30], while the other proposing that wings originated from lateral extension of the notum [31]. A recent study, based on gene expression patterns [32], provided compelling evidence that wings evolved by merging two parts – exites and lateral portion of notum – in the most common ancestor of winged insects (Fig. 1B). Thus, this study gives an insight into how integration of two unrelated GRN modules functioning in different body parts can yield a novel structure.

### 3. Module rewiring (Fig. 1C)

Rewiring of an existing GRN module occurs through reorganization of regulatory linkages among genes. This process can be enhanced by recruitment of novel genes into the existing module and can result in a shift of the module’s developmental role [33, 34]. An example of module rewiring can be seen in the monophyletic clade which includes the two evolutionarily divergent subclasses of sea urchins (Euechinoidea and Cidaroidea) whose skeletogenesis is drastically different embryologically [34]. The gene subcircuits and regulatory linkages that are present in one subclass are missing or operate differently in the other, thus suggesting a dramatic reorganization of the same ancestral GRN at the early stage of sea urchin evolution [34]. Another example of rewiring refers to the *Hox* genes that control development of body plans across many metazoan lineages (Fig. 1C). The reorganization of modules within the *Hox* cluster, at early stages of metazoan evolution, generated the stunning diversity of animal body plans [35–37].

### 4. Module inactivation (Fig. 1D)

Trait loss during evolution is deemed to be caused by mutation(s) in upstream regulatory modules that inactive realization of downstream modules responsible for trait identity [38–40]. For example, Kvon et al. [41] demonstrated that snakes lost their legs due to a few mutations in ZPA enhancer (i.e. Zone of Polarizing Activity Regulatory Sequence), which prevents proper binding of *Sonichedgehog* transcription-factor and hence the realization of downstream limb GRN (Fig. 1D).

The listed examples show that the GRN evolution occurs by modifying one or a few pre-existing GRN modules. The possibility for *de novo* emergence of a GRN that recruits all regulatory elements from different pre-existing GRNs remains possible, however this scenario is less likely as such GRN would require a significantly long evolutionary time to achieve sufficient functionality [42]. To summarize, the aforementioned mechanisms of GRN module evolution fall into three basic categories (Fig. 1E): (1) module birth is birth of a new functional GRN module that occurs by means of the module integration or co-option; (2) module transition is a transformation of pre-existing module from one state to another by the re-wiring mechanism; and (3) module death is the inactivation of a GRN module that eliminates it from the future evolutionary process, resulting in the GRN’s death. Let us assess these mechanisms of module evolution in the light of character and character state concepts.

### GRN evolution supports the invariant nature of a character and state

In an organism GRN modules are nested within each other which imposes a hierarchical structure on the global organismal GRN: during embryo development, the expression of an upstream module acts as regulatory hub that controls the expression of downstream modules, which in turn regulate modules further downstream [29, 34, 43–45]. The entire organismal GRN is a scale-free network, meaning that any smaller module of the network has a similar structure to the whole network but controls fewer processes [14, 46]. Since the global GRN structure is scale-free, the categories of the GRN module evolution are scale-dependent. For example, co-option of a GRN module into a novel body place resulting in emergence of a novel trait represents birth of the new module at this local scale, while at the scale of the entire organism this is a transformation of the global organismal GRN i.e., module transition. On the other hand, the rewiring of a GRN, as in the case of the *Hox* genes cluster (see above), is a module transition at the scale of this cluster, while at the smaller scale, it can be viewed as the birth and death of gene subclusters [47]. So, the limits between the aforementioned mechanisms are somewhat arbitrary and it is logical to believe that GRN evolution occurs under a mixture of the listed mechanisms, especially at the global organismal and time scales. The views that distinguish between character and characters states generally assume that a character is an entity formed by the birth-death process while character states are transformations occurring along the character’s life [2, 11, 12]. I will show that a set of several birth-death events can be seen as both several characters as well as one character, thus eliminating the distinction between the two. This elimination is achieved by forming new states using the combinations of states from several birth-death events that is provided by the modular and scale-free structure of the GRN.

Suppose there is a hypothetical species whose entire GRN consists of five GRN modules {*α, β*_1_, *β*_2_, *γ*_1_, *γ*_2_} (Fig. 2A-B). The evolution of the GRN modules in this species occurs by the birth and death of the modules {*β*_1_, *β*_2_, *γ*_1_, *γ*_2_} in accordance with their hierarchical dependencies: the module *α* is the base module that is always present; the module *β*_2_ is nested in the module *β*_1_, and the module *γ*_2_ is nested in the module *γ*_1_. The first approach (Fig. 2C) to code the module evolution is to use four two-state characters whose character states represent the presence/absence of the modules {*β*_1_, *β*_2_, *γ*_1_, *γ*_2_}. The dependencies within *β* and *γ* module pairs (e.g., presence of *β*_2_depends on the presence of *β*_1_) must be taken into account to impose dependencies between their corresponding characters. The second approach (Fig. 2D) to formalize evolution is to merge the dependent pairs of characters – *{*#1, #2*}* and *{*#3, #4*}* – from the first approach, which yields two three-state characters (Fig. 2D). This merging is done by constructing the combinations: the states of a new character are created out of all possible combinations of states from the individual characters. In this case, each state of the newly formed character bears knowledge of the presence/absence of the two modules simultaneously. The combinations of the modules introduce transformational direction between the states within each character (Fig. 2D). The third approach (Fig. 2E) implies further construction of the combinations by merging the two characters from the second approach. This merging results in one character with nine states: each state contains information on the presence/absence of the four modules simultaneously. It is noteworthy that not all transitions are possible among the nine states. The transitions are structured in a way allowing only birth or death of a single module for each evolutionary step. So, the construction of the combinations introduces a particular penalty (“distance”) between the character states within the character, meaning that the transition to the future state given the present state implies the minimal number of changes. This is biologically logical as, on average, longer time is required to achieve greater evolutionary divergence. Therefore, the birth and death of several GRN modules can be equally represented using a set of a few characters or one character with the appropriate penalty between the states. The representation of module evolution using the first approach requires incorporation of correlation between characters, while the combinations of states (the second and third approaches) substitutes this correlation with the penalty between the states. The approach of constructing the combinations and thereby combining several characters can be extrapolated to formalize evolution of any arbitrary GRN. This indicates that regardless of the scale at which observations are taken – single GRN modules, group of modules or all modules of an organism – the GRN evolution can be equally represented using a set of characters or just one character. Thus, at the level of GRNs, characters and character states are endowed with an invariant property with respect to each other.

**Figure 2.**
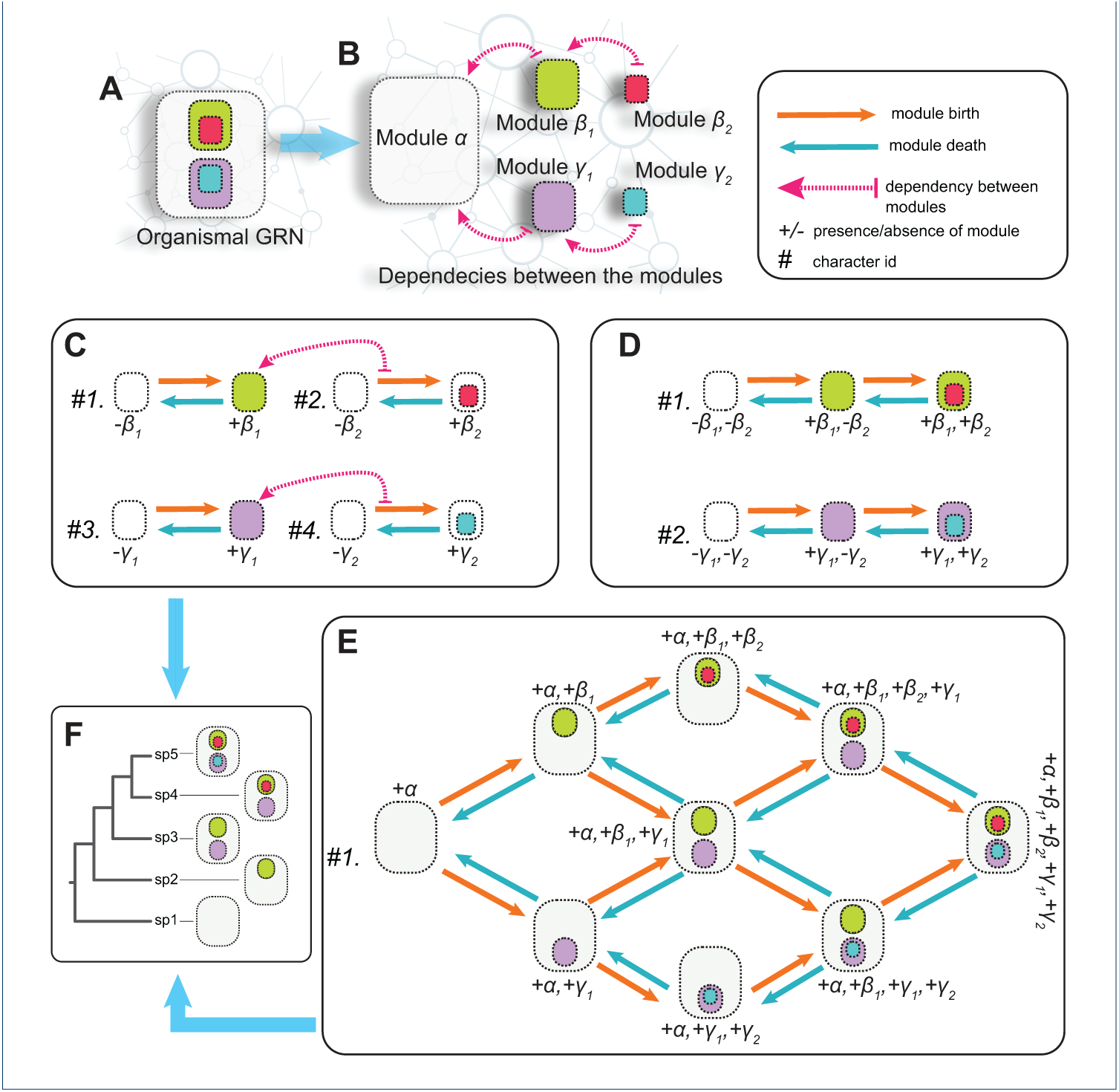
Evolution of GRN modules and construction of the combinations of states. A hypothetical organism **(A)** composed of five GRN modules {*α, β*_1_, *β*_2_, *γ*_1_, *γ*_2_} that are hierarchically dependent **(B)**: *α* is constant, *β*_2_ is nested within *β*_1_, and *γ*_2_ is nested within *γ*_1_. The evolution of the GRN occurs by birth and death of the modules {*β*_1_, *β*_2_, *γ*_1_, *γ*_2_} and can be represented using three coding approaches. **(C)** The first approach uses four two-state characters; this approach requires incorporation of the dependencies (shown with the dashed arrow). **(D)** The second approach uses two three-state characters. **(E)** The third approach uses one character with nine states (absent modules are not shown in the letter notations). The transitions in the second and third approaches are penalized to allow only a birth or death of a single module over one evolutionary step. The state space in the second and third approaches is combinations of the states from the first approach. **(F)** Imaginary species used for demonstrative inference. The tree inference using the set of binary characters from **(C)** and one multistate character with penalties from **(E)** produces that same expected phylogenetic tree shown here.

Morphological traits are the product of GRN modules that makes them have moular and hierarchical properties [48] similar to those of the GRN. For example, the hierarchical dependencies between the organismal body parts and traits associated with them reflect sequential realization of the GRN modules during the embryo development. In addition, trait evolution at the global scale is always associated with birth and loss (death) of traits (see also the section “Mechanisms of GRN evolution”). This means that, likewise GRN modules, phenotypic traits can be also described using a set of characters or just one complex character derived through state combinations and equipped with the appropriate penalties. Moreover, merging several separate characters into one single character can naturally incorporate dependencies if they exist between the traits. I do not suggest that all individual characters have to be combined into one for a phylogenetic analysis - this will make the inference computationally infeasible given present methods. However, the invariant property may be used to decrease the subjectivity associated with character coding as it does not require *a priori* classification of morphological features into characters and character states. Hence, this property is useful for the cases where separation between character and state is ambiguous or morphological structures are known to be dependent.

In practice, constructing states from the combinations of the initial characters (i.e. merging characters into one character) requires selection of an adequate penalty for the state transitions. In phylogenetic inference with “model-based” methods (maximum likelihood or Bayesian), this penalty can be naturally incorporated by appropriately structuring the rate matrix of Markov models. In this case, the penalty is expressed by the infinitesimal rates in the transition rate matrix, and the rate parameters can be tuned so to reflect either dependent or independent evolution between the traits [5, 49, 50]. To demonstrate the effect of penalty, I performed the tree inference for the imaginary species from Fig. 2F in Bayesian framework using *RevBayes* [51] and *Mk*-like models [52] for trait evolution (see also Supplementary Materials). These species have different combinations of GRN modules and their expected phylogenetic tree is the one that reflects a sequence of module gain during the evolutionary course (i.e., the tree in Fig. 2F). The tree inference when GRN modules were coded as four binary characters (i.e., the approach in Fig. 2C), as well as when these four characters were amalgamated into one multistate character with penalties (i.e., the approach in Fig. 2E) converged on the expected tree. However, if the amalgamated multistate character is used without penalties (all transitions between modules are possible) then the inference yields a completely unresolved topology (Supplementary Materials). Thus, the correct structuring of rate matrix is important to maintain the invariant property. The use of the invariant property in parsimony-based methods is, by far, problematic due to the lack of the appropriate approach. The cost-matrix can be potentially used to construct the penalties (reviewed in [13]). However, the use of the cost-matrix in the present context requires additional research.

## Conclusions

This study shows that, in a phylogenetic context, modular and scale-free structure of GRN eliminates distinction between character and character state making them invariant with respect to each other. Since morphological traits are the products of GRNs, this invariance also allows encoding the traits and phenotypes using a set of characters or just one character with multiple states. The invariant property does not require *a priori* classification of morphological features into characters and states, and thereby can be used to reduce subjectivity associated with character coding. Here, I use the demonstrative example to show how the invariant property can be implemented in practice using software for phylogenetic inference.

## ACKNOWLEDGEMENTS

This work was conducted while a Postdoctoral Fellow at the National Institute for Mathematical and Biological Synthesis, an Institute sponsored by the National Science Foundation through NSF Award #DBI-1300426, with additional support from The University of Tennessee, Knoxville. I am grateful to Brian O’Meara (University of Tennessee), Sarah Flanagan, Nicholas Panchy (National Institute for Mathematical and Biological Synthesis, University of Tennessee), Dimitar Dimitrov (University of Copenhagen, Natural History Museum of Denmark) and Ken Puliafico (USDA US Forest) for useful suggestions on the text of the manuscript.

## Supplementary Materials

The datasets supporting the conclusions of this article are included within the article (and its additional files).

